# GenoGAM: Genome-wide generalized additive models for ChIP-seq analysis

**DOI:** 10.1101/047464

**Authors:** Georg Stricker, Alexander Engelhardt, Daniel Schulz, Matthias Schmid, Achim Tresch, Julien Gagneur

**Affiliations:** Gene Center and Department of Biochemistry, Ludwig-Maximilians-Universität München, Feodor-Lynen-Strasse 25, 80333 Munich, Germany.; Technische Universität München, Department of Informatics, Boltzmannstr. 3, 85748 Garching, Germany.; Institut für Medizinische Biometrie, Informatik und Epidemiologie, University Hospital Bonn, Sigmund-Freud-Strasse 25, 53105, Bonn, Germany.; Institute for Genetics, University of Cologne, Zülpicher Str. 47b, 50647 Cologne, Germany.

## Abstract

**Motivation:** Chromatin immunoprecipitation followed by deep sequencing (ChIP-Seq) is a widely used approach to study protein-DNA interactions. Often, the quantities of interest are the differential occupancies relative to controls, between genetic backgrounds, treatments, or combinations thereof. Current methods for differential occupancy of ChIP-seq data rely however on binning or sliding window techniques, for which the choice of the window and bin sizes are subjective.

**Results:** Here, we present GenoGAM (Genome-wide Generalized Additive Model), which brings the well-established and flexible generalized additive models framework to genomic applications using a data parallelism strategy. We model ChIP-Seq read count frequencies as products of smooth functions along chromosomes. Smoothing parameters are objectively estimated from the data by cross-validation, eliminating ad-hoc binning and windowing needed by current approaches. GenoGAM provides base-level and region-level significance testing for full factorial designs. Application to a ChIP-Seq dataset in yeast showed increased sensitivity over existing differential occupancy methods while controlling for type I error rate. By analyzing a set of DNA methylation data and illustrating an extension to a peak caller, we further demonstrate the potential of GenoGAM as a generic statistical modeling tool for genome-wide assays.

**Availability:** Software is available from Bioconductor: https://www.bioconductor.org/packages/release/bioc/html/GenoGAM.html

**Supplementary information:** Supplementary information is available at *Bioinformatics* online.

## Introduction

Chromatin immunoprecipitation followed by deep sequencing (ChIP-Seq) is the reference method used for genome-wide quantification of protein-DNA interactions (Robertson *et al.*, 2007). It is used to study a wide range of fundamental processes covering transcription, replication, and genome maintenance.

ChIP-Seq consists of cross-linking DNA with chromatin, followed by DNA fragmentation and immunoprecipitation of the protein of interest along with its bound DNA fragments. The DNA fragments are then released, amplified, and sequenced. ChIP-Seq has been applied to study DNA-bound proteins of various functions and therefore with various patterns of distribution along the genome. These include transcription factors that are bound at discrete binding sites (Johnson *et al.*, 2007; Barski *et al.*, 2007), histone modifications (Albert *et al.*, 2007; Barski *et al.*, 2007) which are found at nucleosomes, or the transcription (Barski *et al.*, 2007) and replication machinery which are even more broadly distributed.

Often, the quantities of interest are the occupancies relative to technical controls such as the input (a sample that was not subject to the immunoprecipitation step), between genetic backgrounds, treatments, or combinations thereof. Testing for local ocupancy differences between multiple groups of ChIP-Seq samples is therefore an important goal in ChIP-Seq analysis.

In principle, testing could be performed at every single base of the genome. However, the per-base coverage is in practice too low and too noisy for such an approach to have enough statistical power. Testing for differential occupancy has been therefore done by integration of data over larger regions. Generally, benchmarks have shown that differential occupancy perform better if they handle replicate samples (Steinhauser *et al.*, 2016). DESeq tests for differential overall occupancies at pre-defined regions of interest by testing for differences in number of reads overlapping the region (Anders and Huber, 2010). Complementary to testing for overall occupancies, MMdiff (Schweikert *et al.*, 2013) allows testing for differences in shapes in given regions. Other approaches are scanning the genomes or large areas for local differential occupancies. These include diffReps (Shen *et al.*, 2013), where a sliding window moves along the genome in a fixed step size and a robust test based on negative binomial distribution is performed on the number of reads falling into the window. PePr (Zhang *et al.*, 2014) follows a similar scanning approach and estimates local variance. Lun et al. (Lun and Smyth, 2014) devises how to test differential occupancies across windows in given regions while properly controlling for false discovery rate. Their approach has been published in an R package called `csaw`. THOR (Allhoff *et al.*, 2016) uses a Hidden Markov Model approach to segment the genome into regions that are enriched, depleted or not differentially occupied. Of these methods, only DESeq and csaw, which are based on generalized linear models (Nelder and Wedderburn, 1972), can go beyond comparisons between two groups of samples by supporting any full factorial designs including crossed designs. Moreover, all current methods rely on binning or sliding window techniques, for which the choice of the window and bin sizes are not data-driven but subjective.

Here, we introduce GenoGAM, which brings generalized additive models to genomic applications (Fig. 1). Generalized additive models are extensions of generalized linear models for which covariates can be modeled as smooth functions (Hastie *et al.*, 1986). We use them to model ChIP-Seq count rates along the genome. GenoGAM normalizes for sequencing depth and can handle factorial experimental designs, including biological replicates and multiple controls. The amount of smoothing is estimated in an automatic, data-driven manner and thus avoids introducing subjectivity from the analyst. When analyzing differential binding in a factorial design, we obtain well-calibrated per-base-pair p-values and region-wise p-values. Application to a dataset of yeast shows that GenoGAM is more sensitive than state-of-the art differential occupancy methods.

**Fig. 1.**
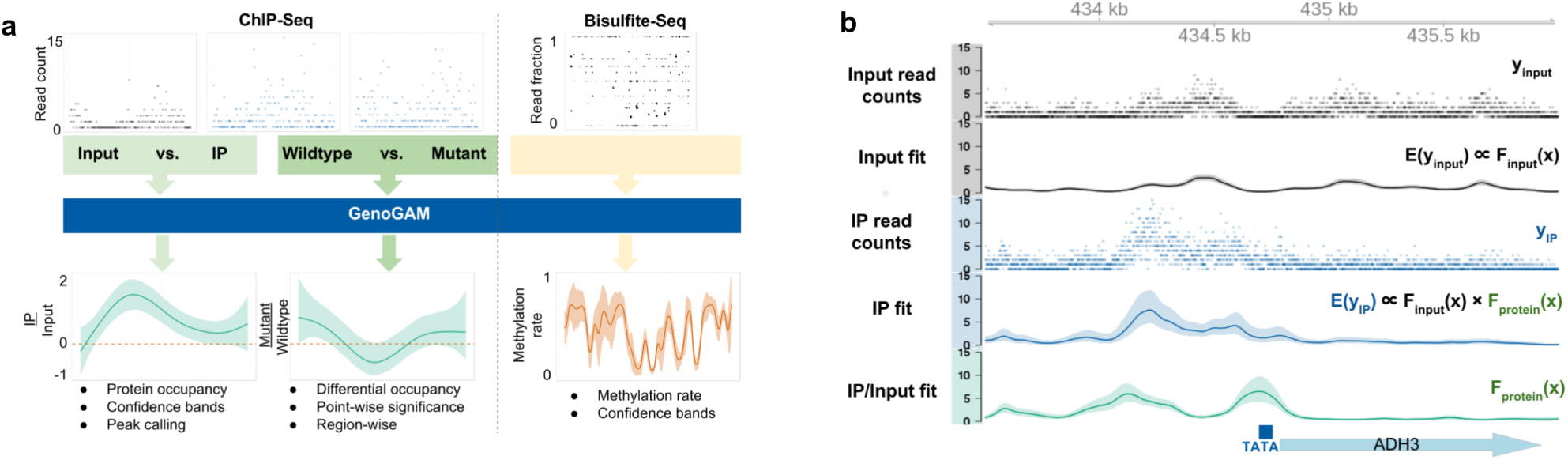
GenoGAM applications and concept. (a) GenoGAM provides a general framework to analyze ChIP-Seq data for both absolute (left arrow) and differential protein (center arrow) occupancy. It can also be applied to infer DNA methylation rate from bisulfite sequencing data (right arrow). (b) ChIP-Seq analysis with GenoGAM yields base-pair resolution occupancy profiles with confidence bands. Input (black) and IP (blue) centered read counts (dots) and fitted smooth (solid line) with 95% confidence intervals (ribbons) for the transcription factor TFIIB for a section of the chromosome XIII of S. cerevisiae. Additionally, the extracted fold change of IP over Input (green) and gene annotation at the very bottom. Simplified equations depict model constituents.

GenoGAM brings further modeling flexibility for which we provide proof-of-principle applications. As a generalized additive model, GenoGAM offers flexible choice of the response distribution. Using proportions rather than absolute counts as response variables, we show that GenoGAM is applicable to estimate DNA methylation rate from bisulfite sequencing data. Moreover, the smooth functions fitted by GenoGAM can be analyzed analytically. Our results indicate that this can be used to identify summits of narrow ChiP-Seq peaks with accuracy close to state-of-the art peak callers.

## Methods

### A generalized additive model for ChIP-Seq data

We consider an experiment consisting of a set of ChIP-Seq samples. A data point is defined by a pair of a ChIP-Seq sample and a genomic position. We denote by *x*_*i*_ the genomic position of the *i*-th data point, by *j*_*i*_ its ChIP-Seq sample and by *y*_*i*_ ≥ 0 the number of fragments in sample *j*_*i*_ centered at position *x*_*i*_. For single-end libraries, the fragment center is estimated by shifting the read end position by a constant. In case of single end data, the fragment length *d* was estimated using the Bioconductor package `chipseq` and its *coverage* method. It is defined as the optimal shift for which the number of bases covered by any read is minimized. Thus, the center was taken as the start of the read shifted by 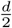 downstream. When reducing ChIP-Seq data to fragment centers rather than full base coverage, each fragment is counted only once. This reduces artificial correlation between adjacent nucleotides.

We model the counts *y*_*i*_ using the following generalized additive model:

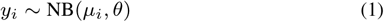

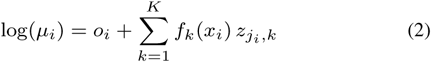

The counts *y*_*i*_ are assumed to follow a negative binomial distribution with means *µ*_*i*_ (Equation 1) and a dispersion parameter *θ* that relates the variance to the mean such that 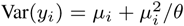. Consequently, the model accounts for dispersion beyond Poisson noise (Anders and Huber, 2010).

The logarithm of the mean *µ*_*i*_ is the sum of an offset *o*_*i*_ and one or more smooth functions *f*_*k*_ (Equation 2). The offsets *o*_*i*_ are predefined data-point specific constants that account for sequencing depth variations (see subsection Sequencing depth variations). The indicator variable *z*_*j*_*i*,__*k* is 1 if the smooth function *f*_*k*_ contributes to the mean counts of sample *j*_*i*_ and 0 otherwise. As shown below, this formulation allows modeling IP versus input experiments as well as factorial experimental designs.

We model IP versus input experiments using GenoGAM with two smooth functions: *f*_input_ that contributes to both input and IP samples, and *f*_protein_ that only contributes to IP samples. More specifically, *f*_input_ models local ChIP-Seq biases common to input and IP, whereas *f*_protein_ models the protein log-occupancy up to one genome-wide scaling factor. Figure 1b shows the application of this model to one ChIP-Seq library for the *S. cerevisiae* general transcription factor TFIIB and its input control (Supplementary Methods).

In GenoGAM, the smooth functions are represented by cubic spline curves, which are written as linear combinations of a set of regularly spaced basis functions *b*_*r*_, i.e. 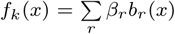. We chose second order B-splines as basis functions, which are bell-shaped cubic polynomials over a finite support (De Boor, 1978). To avoid overfitting, regularization of the functions *f*_*k*_ is carried out by penalization of the second order differences of the spline coefficients, which approximately penalizes second order derivatives of *f*_*k*_ – an approach called P-splines or penalized B-splines (Eilers and Marx, 1996). The optimization criterion for P-splines is the sum of the negative binomial log-likelihood (depending on the response vector *y* and the vector *β* containing the coefficients of all smooth functions) plus a penalty function that is weighted by the smoothing parameter *λ*:

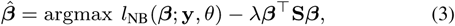

where S is a symmetric positive matrix that encodes the squared second order differences of the coefficients *β* (Eilers and Marx, 1996). This regularization allows dense placements of the basis functions (between 20 and 50 bp), while relying on the smoothing parameter *λ* to protect against overfitting. Large values of *λ* yield smoother functions. A single smoothing parameter common to all smooth functions proved to be sufficient for our applications. For given *λ* and *θ*, model fitting was performed using penalized iteratively re-weighted least squares (See subsection Model fitting).

Adapting a Bayesian view, the penalized likelihood can be interpreted as a posterior probability, and the penalization term arises from a Gaussian prior on the coefficients *β*. Large-sample approximations then yield a multivariate Gaussian posterior distribution for *β*, and, by the linearity of 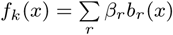, Gaussian posteriors for the point estimates *f*_*k*_(*x*). This allows for the construction of pointwise confidence bands (Wood, 2006). An example of the fitted smooth functions and their confidence bands for the yeast transcription factor TFIIB is shown in Figure 1b.

### Fitting of a GAM on a genome-wide scale, given the smoothing and dispersion parameters *λ* and *θ*

Since the computation time of a GAM grows polynomially with the number of basis functions, fitting one model to a whole chromosome is unfeasible. Instead, we propose to fit separate GAMs on sequential overlapping intervals (or tiles, Supplementary Fig. S1a). Each chromosome was partitioned into equally-sized intervals called chunks. Tiles were defined as chunks extended on either side by equally-sized overhangs. The generalized additive model was fitted on each tile separately using the *gam* function of the R package `mgcv`. Point estimates at each base pair of the smooth functions and their standard errors were extracted with the *predict* function on the fitted object setting “type” parameter to “iterms”. The tile fits were then restricted to their chunk to define the chromosome-wide fit. As overlap length increases, agreement of the fit at the midpoint of the overlapping region increases. A genome-wide fit is obtained by joining together tile fits at overlap midpoints (Supplementary Fig. S1a). This approximation yields computation times that are linear in the number of basis functions at no practical precision cost (Fig. S1b). Furthermore, it allows for parallelization, with speed-ups being linear in the number of cores (Supplementary Fig. S1c). This approximation parallelizes the computation over the data, which will allow future implementation of GenoGAM in map-reduce frameworks such as Spark (Zaharia *et al.*, 2010).

### Data-driven determination of the smoothing and dispersion parameters *λ* and *θ*

To determine the optimal value for *λ* and *θ*, we tried generalized cross-validation, an analytical leave-one-out large-sample approximation (Wood, 2006). However, this yielded very wiggly fits indicative of overfitting. We thus developed an empirical cross-validation scheme.

For efficiency, cross validation was performed using only a subset of the data. We selected a sufficiently large set of distinct regions that are long enough to not suffer from border effects common to spline fitting. Using 20 or more distinct regions containing at least 100 basis functions gave satisfactory empirical results (Supplementary Table S1). For peak calling purposes, regions were selected that had the most significant fold change of IP versus input read counts.

In each region, 10-fold cross-validation was performed, where a tenth of the data points were removed, the model was fitted on the remaining data points, and the log-likelihood of the left-out data points was computed. To avoid overfitting due to short range correlations, each cross-validation fold did not consist of randomly selected single genomic positions, which would recapitulate the leave-one-out scheme, but of short intervals. The length of these intervals was was set to 20 bp (approximately a tenth of the fragment length.) in absence of replicates and twice the average fragment sizes when replicates were available.

For a given pair of values for *λ* and *θ*, the score function was defined as the sum of out-of-sample log-likelihood over all cross-validation folds and all tiles, restricted to the data points within chunks to not depend on poor fitting in overhangs. Investigation on grid values of *θ* and *λ* showed that the out-of-sample log-likelihood was typically unimodal. We therefore used a numerical optimizer to jointly fit the two parameters (R function *optim*, BFGS method with default finite-difference approximation of the gradient) (Broyden (1970), Fletcher (1970), Goldfarb (1970), Shanno (1970)).

### Sequencing depth variations

We used an approach originally suggested by Meyer and Liu (2014) that was robust to variations in signal-to-noise ratio. Variations for sequencing depth was controlled by using size factors computed by DESeq2 (Love *et al.* (2014) version 2_1.10.0). This method robustly estimates fold-changes in overall sequencing depth by comparing read counts of predefined regions. The selection criteria for these regions was application-specific. For differential binding application, all tiles were considered. For IP versus input comparisons (TFIIB and peak calling applications see Supplementary information), the selected regions were the 1,000 tiles with smallest p-value according to DESeq2 test for enrichment of IP over input performed on total read counts per tile. This allowed to select tiles that were most likely containing peaks.

### TFIIB dataset

This dataset consisted of two samples: one input and one IP without replicates (Processing in Supplementary information). Hence, there was no need for an offset. We used the following GenoGAM model:

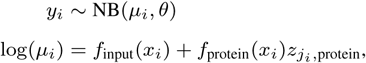

where *z*_*j*_*i*,_protein_ = 1 whenever *j*_*i*_ is the index of an IP sample and *z*_*j*_*i*,_protein_ = 0 whenever *j*_*i*_ is the index of an input sample. Further parameter details are given in Supplementary Table S1.

### Differential binding

#### Data

This dataset consisted of four samples: two biological replicate IPs for the wild type strain and two biological replicate IPs for the mutant strain. Data processing and gene boundaries definition are described in Supplementary information.

#### GenoGAM model

We used a GenoGAM model that compares the mutant with the wildtype ChIP data as follows:

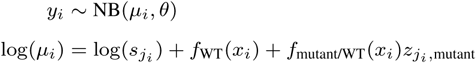

where *z*_*j*_*i*_,mutant_ = 1 for *j* index of mutant samples and 0 for wild-type samples. The offsets log(*s*_*j*_*i*__) are log-size factors computed to control for sequencing depth variation and overall H3K4me3 across all four samples (see ‘Sequencing depth variations’ above).

#### Position-level significance testing

Null hypotheses of the form *H*_0_: *f*_*k*_(*x*) = 0 for a smooth function *f*_*k*_() at a given position *x* of interest were tested assuming approximate normal distribution of the corresponding z-score, i.e.:

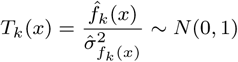

where 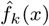 and 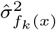 denote point estimate and standard error of the smoothed value using the predict function of the R package mgcv (Wood, 2006).

#### False discovery rate for predefined regions

Let *R*_1_,…, *R*_*p*_ be *p* regions of interest, where a region is defined as a set of genomic positions. Regions are typically, but not necessarily, intervals (e.g. genes or promoters). For instance, all exons of a gene could make up a single region. Regions can be a priori defined or defined on the data using independent filtering (Bourgon *et al.*, 2010). For instance, when testing for significant differences between two conditions, regions can be selected for having a large total number of reads over the two conditions (Lun and Smyth, 2014).

For *j* in 1,.., *p*, let 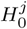 be the composite null hypothesis that the smooth function *f*_*k*_ values 0 at every position of the region *R*_*j*_. The False Discovery Rate was controlled as in Lun and Smyth (2014):

1. Position-level p-values at all region positions were computed using position-level significant testing as described above.
2. Within each region *R*_*j*_, position-level p-values were corrected for multiple testing using Hochberg family-wise error rate correction (Hochberg, 1988). The p-value for the null hypothesis 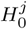 was then computed as the minimal family-wise error rate corrected position-level p-value. This step gives one p-value per region.
3. FDRs were controlled using the Benjamini-Hochberg procedure (Benjamini and Hochberg, 1995) applied to the region-level p-values.

#### Benchmarking

THOR allows to restrict the search to pre-defined regions. However, running THOR in this mode yielded worse results. We therefore called differential binding sites with a genome-wide run of THOR and then overlapped them with the gene coordinates. The same was done for PePr and diffReps, because these methods do not allow for a fixed set of regions as input. Supplementary information gives further details on how the competitor methods were run.

Gene expression levels were computed as the median normalized probe levels for the three replicate YPD conditions of all tiling array probes provided by Xu *et al.* (2009) overlapping gene coordinates defined by Xu *et al.* (2009). Genes from Xu *et al.* (2009) and from Thornton *et al.* (2014) were matched by symbol.

To compute ROC curves, binary labels (expressed = 1 if above a given expression level quantile cutoff, or not = 0 otherwise) were assigned to each gene, and for each method, genes were ranked according to their respective significance value. For THOR, PePr and diffReps, genes that did not overlap any differentially bound site, p-values were set to 1. Then, ROC curves and AUC for all expression level quantile cutoffs in steps of 0.01 were computed.

### Methylation data

To model *y*_*i*_, the number of reads of methylated state, out of *n*_*i*_, the total number of reads, we used the quasi-binomial model defined by:

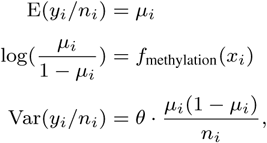

where the scale parameter *θ* > 0 models dispersion. The model was applied on only one tile with a width of 120 kb, reproducing Figure 2A of Smallwood et al. (Smallwood *et al.*, 2014) (Fig. 4). Further parameter details are given in Supplementary Table S1.

**Fig. 2.**
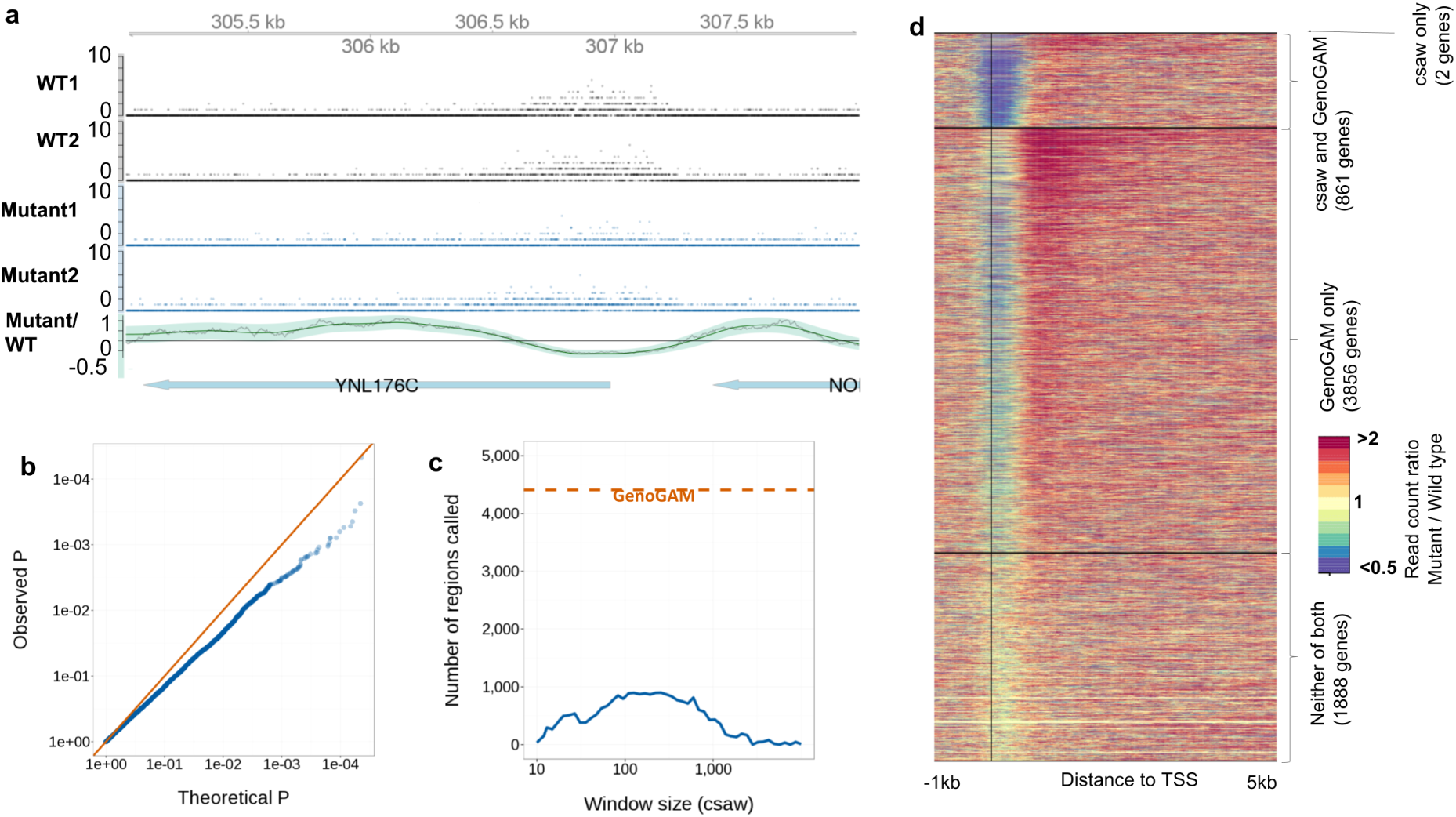
Statistical testing for factorial designs. (a) Read counts (dots) and fitted rates with 95% confidence bands for wild-type (black) and mutant (blue) and the log-ratio of mutant over wild-type with confidence band (bottom row, green) around YNL176C. For comparison, log-ratios computed in sliding windows of size 184bp (bottom row, gray, optimized window size, see section ‘Comparison of GenoGAM fit with sliding window smoothing’). (b) Empirical (y-axis) versus theoretical (x-axis) p-values in base-level permuted count data (Supplementary Methods). P-values at every 200 bp positions are shown. (c) Number of genes with significant differential occupancies in mutant over wild type (FDR < 0.1) reported by GenoGAM (orange) and by csaw (blue) as function of window size (x-axis). (d) Fold-change of counts in mutant over wild-type in 150 bp windows for all 6607 yeast genes in the -1 to 5 kb region centered on TSS (vertical black line). The genes are sorted into four groups (separated by the black horizontal lines) according to which method reports them significant. From top to bottom: csaw only (2 genes), csaw and GenoGAM (861 genes), GenoGAM only (3,856 genes) and none (1888 genes). Within each group genes are ordered by p-values (lowest to highest from top to bottom). The “csaw and GenoGAM” group is sorted by GenoGAM p-values. Comparisons to all other methods can be found in Supplementary Figure S3 - S5

**Fig. 3.**
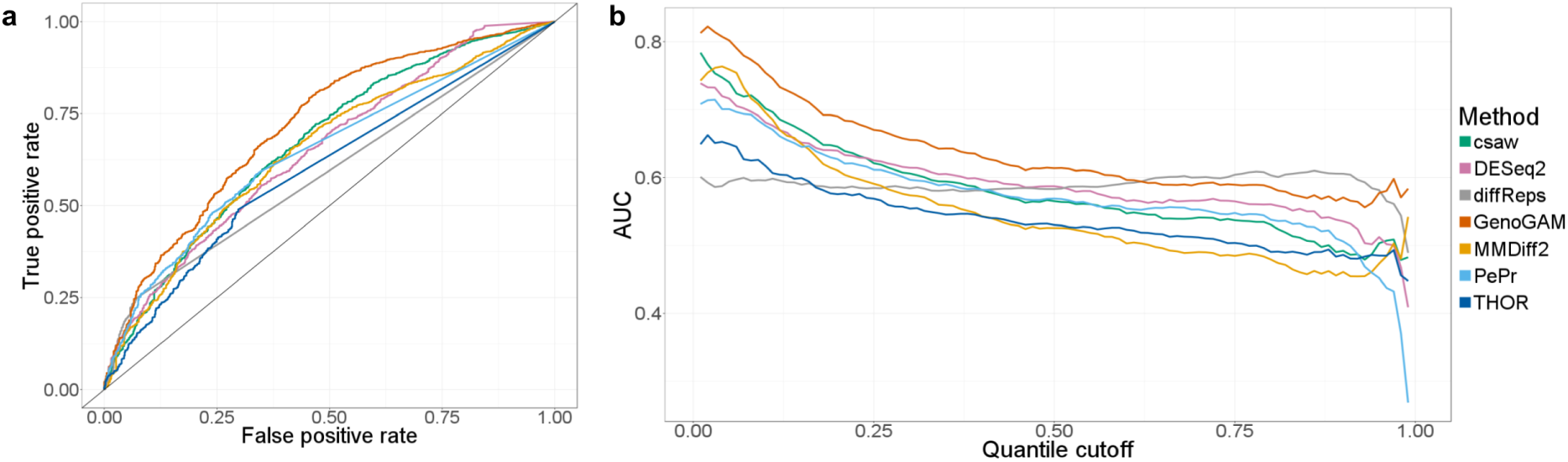
GenoGAM identifies differential regions with greater Recall. (a) Area under the curve (AUC) for all possible quantile cutoffs from 0 to 1 in steps of 0.01. Up to a cutoff of 0.6, GenoGAM (red) performs consistently better than all competitor methods by around 0.03-0.04 points above the second best method (csaw and DESeq2, green and pink, respectively). The entire range of quantile cutoffs is shown out of completeness, reasonable values are between 0.15 and 0.25. (b) ROC curve based on a quantile cutoff of 0.15 (see Figure S7 for other cutoffs). GenoGAM has a constantly higher recall with a lower false positive rate. The partially straight lines for THOR, PePr and diffReps are stemming from tied genes with no significance value. (c) Boxplots showing gene expression levels for significant and not significant genes according to the respective threshold of the method (0.1 for FDR based methods and 1e-5 for PePr). Gene expression levels are clearly distinct between the group of significant and not significant genes across all methods

**Fig. 4.**
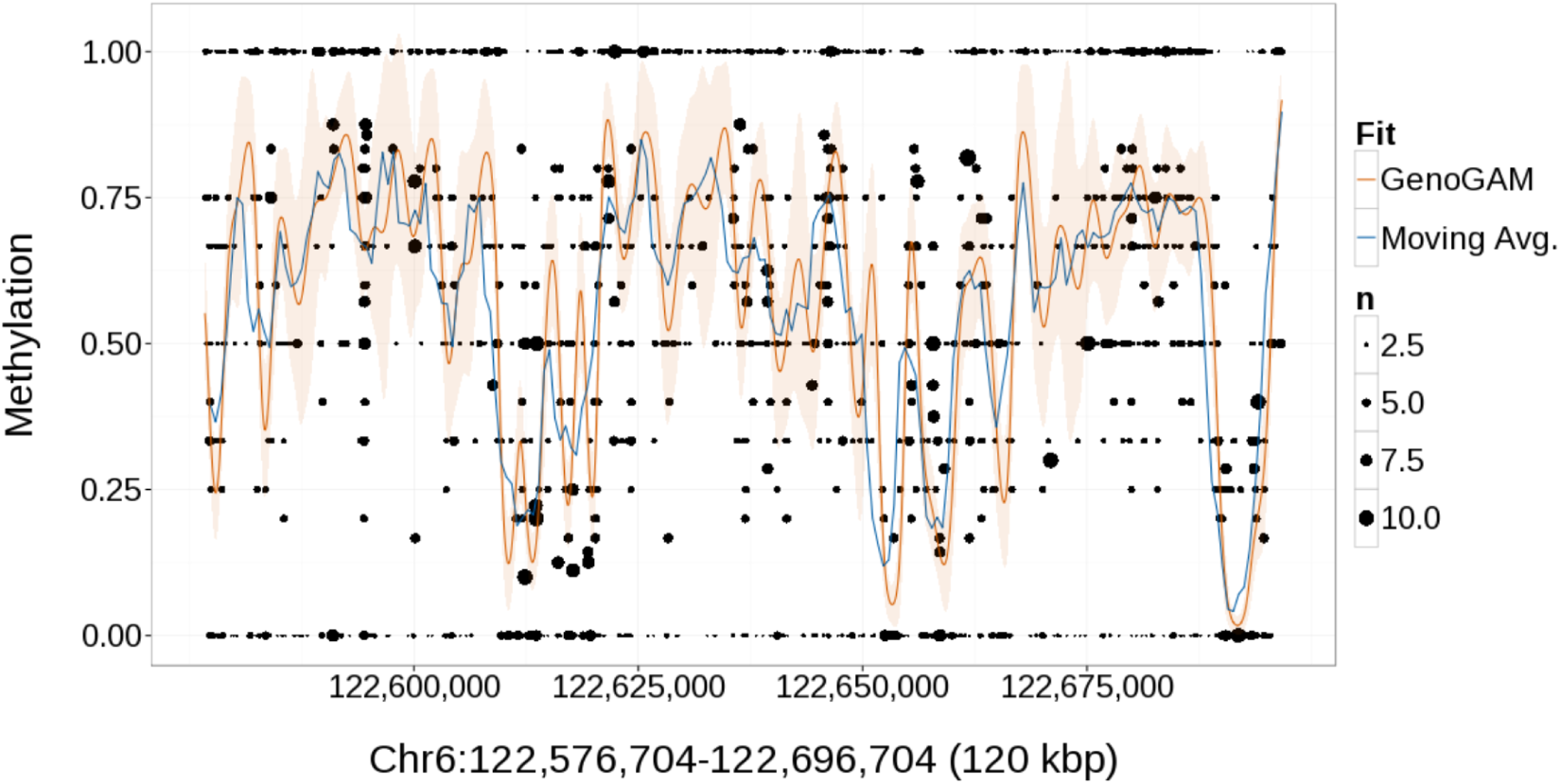
Application to DNA methylation data. Estimated DNA methylation rates in a 120 kb region of chromosome 6 of the mouse (cf. Smallwood et al.Smallwood et al. (2014)). Shown are the data for bulk embryonic mouse stem cells grown in serum; ratios of methylated counts for each CpG position (black dots), with point size proportional to the number of reads. The estimated rates are shown for the moving average approachSmallwood et al. (2014) of 3,000 bp bins in 600 bp steps (blue line) and for the GenoGAM (orange line) with 95% confidence band (ribbon).

## Results

### Testing for differential occupancy with GenoGAM with controlled type I error

To assess the performance of GenoGAM for calling differential occupancy, we re-analyzed histone H3 Lysine 4 trimethylation (H3K4me3) ChIP-Seq data of a study (Thornton *et al.*, 2014) comparing wild type yeast versus a mutant with a truncated form of Set1, the H3 Lysine 4 methylase. H3K4me3 is a hallmark of promoters of actively transcribed genes. Thornton and colleagues (Thornton *et al.*, 2014) have reported genome-wide redistribution in the truncated Set1 mutant of H3K4me3, which is depleted at the promoter and enriched in the gene body. This dataset is interesting for differential occupancy analysis because it is not about the overall number of counts, but about the redistribution of H3K4me3 within the gene. Hence, methods must be sensitive to differences at any location within the gene. We expect such redistribution of the mark at all genes that are transcriptionally active for yeast cells grown in rich media.

We modeled the two replicate IPs for mutant and for the wild type with GenoGAM using one smooth function *f*_WT_ for the wild type reference occupancy, and one further smooth function *f*_mutant/WT_ for the differential effect (Methods). The offsets were computed to control for variations in sequencing depth between replicates and overall genome-wide H3K4me3 level (Methods). This yielded base-level log-ratio estimates and their 95% confidence bands genome-wide (Methods, Fig. 2a for data and fit at the gene *YNL176C* consistent with the report of reduced binding at promoter regions).

Confidence bands of GAMs are formally Bayesian credible intervals (Methods). However, previous studies based on simulated data showed that these confidence bands have close to nominal coverage probabilities and can, in practice, be used in place of frequentist confidence intervals (Marra and Wood, 2012). We estimated base-level p-values using the point-wise estimates and standard deviations (Methods). To empirically verify that the p-values were at least conservative, we created a negative control dataset by per-base-pair independent permutation of the counts between the four samples. The offsets were set to 0 and the smoothing and dispersion parameters were estimated again. This non-parametric permutation scheme makes less assumptions than previous simulation studies (Marra and Wood, 2012). Nonetheless, per-base-pair p-values in this negative control experiment were slightly overestimated (Fig. 2b). These results show that GenoGAM can be used to identify individual positions of significant differential occupancies with controlled type I error. Here, correction for multiple testing can either be done using the Benjamini and Hochberg procedure (Benjamini and Hochberg, 1995) or procedures that exploit dependencies between adjacent positions (Wei *et al.*, 2009).

Complementary to de novo identification, predefined regions, such as genes, can be tested for differential occupancies. To test for differences at any position in a region using GenoGAM, we propose to apply Hochberg’s procedure to correct the pointwise p-values for multiple testing, and to report the smallest of these corrected p-values (Methods). We confirmed by permutation analysis that this approached conservatively controlled for type I error rate (Supplementary Fig. S2).

### Higher sensitivity in testing for differential occupancy

We first compared GenoGAM to csaw, which is its most directly comparable method because only GenoGAM and csaw can model flexible factorial designs and assess differences in overall read counts and in shape. One fundamental difference is that csaw is based on a sliding window approach requiring an a priori defined window size. In contrast, the smoothing parameter of GenoGAM is learnt from the data by maximizing the out-of-sample likelihood in cross-validation (Methods). Across all investigated window sizes, the csaw algorithm reported a maximum of 863 significant genes at FDR < 0.1 (Fig. 2c). Moreover, the number of identified genes depended strongly on the choice of the window size (Fig. 2c). In contrast, GenoGAM reported 4,717 significant genes at the same FDR cutoff, which is much closer to the number of transcriptionally active genes (Xu *et al.*, 2009). The genes reported by GenoGAM included all the genes reported by csaw except two, indicating that GenoGAM captured the same signal but with a higher sensitivity (Fig. 2d). The genes reported only by GenoGAM showed a differential occupancy pattern similar yet weaker to the genes common to csaw and GenoGAM, with depletion in the promoter and enrichment in the gene body (Fig. 2d), indicating that GenoGAM captured true biological signal.

We next compared GenoGAM against a comprehensive set of differential occupancy methods that proved to be competitive in a recent benchmark (Steinhauser *et al.*, 2016). These methods were DESeq2 (Anders and Huber, 2010), MMDiff2, the current version of MMDiff (Schweikert *et al.*, 2013), csaw (Lun and Smyth, 2014), diffReps (Shen *et al.*, 2013), and PePr (Zhang *et al.*, 2014). Two more methods highlighted by Steinhauser and colleagues (Steinhauser *et al.*, 2016) were excluded: ChIPComp, as the R package is hardcoded to be used on mouse and human datasets only, and DiffBind, which is redundant, since it is essentially a test for differences in overall counts based on either DESeq2 (already present) or edgeR (used by csaw). We furthermore included the more recently published HMM-based method THOR (Allhoff *et al.*, 2016) (Methods).

The least number of significant genes (FDR < 0.1 or the respective default threshold set by the method) were identified by DESeq2 (735) csaw (863) and diffReps (1193). The most were reported by THOR (2687), PePr (3248) and MMDiff2 (3482), closer to GenoGAM. To make sure that i) the reported genes indeed corresponded to transcriptionnaly active genes (also see Supplementary Fig. S6) and ii) that these results did not depend on FDR cutoffs we performed receiver operating characteristic (ROC) using expressed genes as a proxy for true positives (Methods). GenoGAM had the largest Area Under the ROC curve, when considering that the 15% of the genes with lowest expression levels in Xu *et al.* (2009) are not expressed (Fig. 3a, Methods). Moreover, GenoGAM consistently had the largest AUC for any gene expression cutoff up to 60% genes to be not expressed (Fig. 3b). These results indicate that GenoGAM is more sensitive than current methods for testing differential occcupancy, while still controlling for type I error rate.

### Comparison of GenoGAM fit with sliding window smoothing

In the uncommon situation where a benchmark is available as for the Thornton et al. dataset, one can objectively define an optimal window size for sliding window approaches. The log-ratios estimated by GenoGAM fit well to log-ratios computed in sliding windows of size 184bp, the window size maximizing the area under curve for csaw for a gene expression quantile cutoff of 0.15 (Fig. 2a). Also, the GenoGAM 95% confidence ribbon captures very well the short-range fluctuations of the sliding window estimates. Hence there is a general agreement between the two approaches. However, the benefits of GenoGAM are clear: First, the GenoGAM fit is smooth and differentiable. Second, unlike in the window-based approach, the amount of smoothing is solely estimated from the ChIP-seq data, without prior knowledge from the benchmark.

### Application to DNA methylation data

Generalized additive models are based on the generalized linear modeling framework and thus allow any distribution of the exponential family for the response. Therefore, GenoGAM can be also used to model continuous responses, for instance using the Gaussian distribution, and proportions using the Binomial distribution. For ChIP-Seq data, a log-linear predictor-response relationship of the form (Equation 2) is justified by the fact that effects on the mean are typically multiplicative. However, other monotonic link functions could also be used. Moreover, quasi-likelihood approaches are supported, allowing for the specification of flexible mean-variance relationships (Wedderburn, 1974).

To test the flexibility of GenoGAM, we conducted a proof-of-principle study on modeling bisulfite sequencing of bulk embryonic mouse stem cells grown in serum (Smallwood *et al.*, 2014). Bisulfite sequencing quantifies methylation rate by converting cytosine residues to uracil, leaving 5-methylcytosine residues unaffected. At each cytosine, the data consisted of the number *n*_*i*_ of fragments overlapping the cytosine and the number *y*_*i*_ of these fragments for which the cytosine was not converted to uracil. The quantity of interest was the methylation rate, i.e. the expectation of the ratio *y*_*i*_/*n*_*i*_. In the original publication, single nucleotide position methylation rates were estimated using a sliding window approach with an ad-hoc choice of window size of 3 kb computed in steps of 600 bp. Figure 4 reproduces an original figure showing the fit in a 120kb section of chromosome 6. We modeled this 120 kb section with GenoGAM using a quasi-binomial model, where the response was the number of successes *y*_*i*_ out of *n*_*i*_ trials, the log-odd ratio was modeled as a smooth function of the genomic position, and the variance was equal to a dispersion parameter times the variance of the binomial distribution. Smoothing and dispersion parameters were determined by cross-validation (Methods). The GenoGAM fit was consistent with the original publication (Smallwood *et al.*, 2014), but did not rely on manually set window sizes and provided confidence bands (Fig. 4). As expected, wider confidence bands were obtained in regions of sparse data and tighter bands in regions with a lot of data (Fig. 4).

### Calling ChIP-seq peak summits

The smooth function estimates and their representation as P-splines provided by GAM offer new opportunities for subsequent analyses: First and second order derivatives can be computed immediately. Those can be used to infer summits of ChIP-seq peaks (as positions *x* where *f′*(*x*) = 0 and *f″*(*x*) < 0). Supplementary information shows how a peak caller can be therefore built, including a test for statistical significance of the peaks (Supplementary Figure S8). Comparison of this approach to a few widely used peak callers (MACS (Zhang *et al.*, 2008), JAMM (Ibrahim *et al.*, 2015) and ZINBA (Rashid *et al.*, 2011)) on small size data sets (Human chromosome 22 and yeast) indicates reasonable performance (Supplementary Figures S9 - S12).

### Implementation and current computational limitations

We implemented the method in the freely available Bioconductor R package `GenoGAM`. Given a configuration file of the BAM files, experiment design matrix and model formula, it will automatically estimate all parameters of the model. Alternatively, users can provide their own size factors or smoothing and overdispersion parameters. `GenoGAM` provides downstream analysis functions for differential binding and peak calling as described above. `GenoGAM` supports a number of parallel backends through the Bioconductor parallel framework `BiocParallel`.

The genome-wide analysis on yeast, which is 12 Mb long, presented in this paper took around 20 hours on 60 cores including parameter estimation by cross-validation. Once the smoothing and overdispersion parameters are estimated, the runtime reduces to around an hour. Nonetheless, running time for whole human genome, which is about 3 Gb is long, is at this stage not practical. Our main focus so far has been to establish the framework and to evaluate its statistical properties, which we present in this paper. Solutions to these computational limitations are currently being addressed with promising results, in order to extend GenoGAM to genome-wide application for larger genomes.

## Discussion

We have introduced a generic framework based on generalized additive models to model ChIP-Seq data. Unlike most other methods for ChIP-Seq analysis, GenoGAM is a data generative model, which gives an explicit likelihood of the data. This in turn yields an objective criterion to set the amount of smoothing. Smoothing and dispersion parameters were obtained by cross-validation, i.e. they were fitted for the accuracy in predicting unseen data. This criterion turned out to provide useful values of smoothing and dispersion for inference. Moreover it led to reasonable uncertainty estimates since confidence bands of the fits were found to be only slightly conservative. To our best knowledge, GenoGAM is the first method so far that has addressed the setting of the amount of smoothing for ChIP-Seq data. The possibility exists to estimate the smoothing and dispersion parameters separately for each sample, which would result in more robust estimates at the cost of some flexibility. However, in our analyses the samples within an experiment were all similar enough to estimate the parameters globally.

The utilization of genome-wide GAMs comes with a number of advantages: First, we flexibly model factorial designs, as well as replicates with different sequencing depths using size factors as offsets. More elaborate usage could include position- and sample-specific copy number variations, or GC-biases. Second, applying GAMs yields confidence bands as a measure of local uncertainty for the estimated rates. We showed how these can be the basis to compute point-wise and region-wise p-values. Third, GAMs outputs analytically differentiable smooth functions, allowing flexible downstream analysis. We discussed how peak calling can be elegantly handled by making use of the first and second derivatives. Fourth, various link functions and distributions can be used, providing the possibility to model a wide range of genomic data beyond ChIP-Seq, as we illustrated with a first application on DNA methylation. Hence, we foresee GenoGAM as a generic method for the analysis of genome-wide assays.

Scalability to fit very long longitudinal data such as whole chromosomes at base-pair resolution was made possible by parallelization over the data and allowing approximations rather than exact computation of the fit (Heinis, 2014). Nonetheless, practical usage of our current implementation remains limited to organisms with small genomes such as yeast or bacteria, or to selected subsets of larger genomes, such as promoters. Despite the present limitations in runtime and genome size, we are confident that GenoGAM is of importance to the bioinformatics community. Future research direction includes improving the computing time, for instance leveraging on recent progresses for fitting large GAMs (Wood *et al.*, 2016).

## Acknowledgements

We thank Ulrike Gaul, Ulrich Unnerstall and Michael Lidschreiber for fruitful discussions on data analysis, Martin Morgan and Hervé Pagès for support during the implementation of the GenoGAM package, Stefan Krebs for sequencing and raw data preprocessing, and Ulrich Mansmann and Patrick Cramer for institutional support.

## Author’s contributions

Conceived the project and supervised the work: JG AT. Developed the software and carried out the analysis: GS AE JG Carried out the ChIP-Seq experiments for TFIIB on yeast: DS. Gave advice on statistics: MS. Wrote the manuscript: JG AE GS MS AT

